# EPOC outside the shield: comparing the performance of a consumer-grade EEG device in shielded and unshielded environments

**DOI:** 10.1101/2020.08.16.253229

**Authors:** Jordan Wehrman, Sidsel Sörensen, Peter de Lissa, Nicholas A. Badcock

## Abstract

Low-cost, portable electroencephalographic (EEG) headsets have become commercially available in the last 10 years. One such system, Emotiv’s EPOC, has been modified to allow event-related potential (ERP) research. Because of these innovations, EEG research may become more widely available in non-traditional settings. Although the EPOC has previously been shown to provide data comparable to research-grade equipment and has been used in real-world settings, how EPOC performs without the electrical shielding used in research-grade laboratories is yet to be systematically tested. In the current article we address this gap in the literature by asking participants to perform a simple EEG experiment in shielded and unshielded contexts. The experiment was the observation of human versus wristwatch faces which were either inverted or noninverted. This method elicited the face-sensitive N170 ERP.

In both shielded and unshielded contexts, the N170 amplitude was larger when participants viewed human faces and peaked later when a human face was inverted. More importantly, Bayesian analysis showed no difference in the N170 measured in the shielded and unshielded contexts. Further, the signal recorded in both contexts was highly correlated. The EPOC appears to reliably record EEG signals without a purpose-built electrically-shielded room or laboratory-grade preamplifier.

## 1. Introduction

Electroencephalography (EEG) has traditionally been the province of those who can afford expensive equipment and specialised laboratory spaces. A potential solution to this access problem has been provided by the development of portable and comparatively inexpensive EEG headsets for use in the home (e.g., Emotiv EPOC®, Imec’s wireless EEG headset, NeuroFocus MyndTM, Neurokeeper’s headset, NeuroSky Mindwave®). Though not designed for research, modifications to consumer systems such as the EPOC have been used to timelock EEG data with conditions of interest, allowing analysis of the electrophysiological response to specific events (i.e., event-related potentials, ERPs). In fact, the quality of these recordings has been shown to be equivalent to research-grade EEG systems in a laboratory setting (Badcock et al., 2013; 2015, de Lissa et al., 2015). However, the advantage of these portable systems is the chance to take them outside of laboratory settings. Therefore, especially given the lack of a noise-reducing pre-amplifier used with expensive research-grade EEG systems, it is important to establish the reliability of ERPs recorded inside and outside an electrically shielded room using consumer-grade EEG systems. Here, we specifically considered Emotiv’s EPOC.

Though a comparison of signals recorded between shielded and unshielded environments has yet to be performed, this has not stopped researchers from using the device in non-traditional settings. For example, Debener, Minow, Emkes, Gandras, and de Vos (2012) showed that the EPOC could be used to track ERPs while in a quiet office or walking outdoors (see also De Vos, Gandras, & Debener, 2014). Further, due to its quick setup, the EPOC may be a useful tool in gaining electrophysiological insights from individuals or populations who find the traditionally longer setup time challenging (e.g., individuals with autism; Yau, McArthur, Badcock, & Brock, 2015).

While the EPOC is a tempting surrogate research EEG device, two primary questions must be answered regarding its appropriateness. Firstly, the data retrieved from the EPOC must be comparable to data from research-grade EEG systems. Fortunately, this appears to be the case. De Vos, Kroesen, Emkes, and Debener (2014) demonstrated that the EEG amplifiers used in both the EPOC and a wired research-grade EEG system provide comparable measures of the auditory P300 component. While the De Vos et al. (2014) study used two systems sequentially, Badcock et al. (2013; 2015) simultaneously recorded with EPOC and a research-grade Neuroscan system, finding that late auditory ERPs were similar between the systems. Also using simultaneous recordings, de Lissa et al. (2015) found that EPOC reliably records face-sensitive occipito-temporal N170 peaks (when modified to incorporate an alternative reference system to the standard mastoid sensors), recording both face-sensitivity and face-inversion effects which did not differ from those recorded with a Neuroscan system. Thus, multiple groups have demonstrated that the EPOC is suitable to measure ERP effects in auditory and visual modalities.

While research-grade EEG recording usually takes place in an electrically-shielded room, one advantage of relatively inexpensive consumer-grade EEG headsets such as the EPOC system is its portability and ease of set-up. For example, de Wit et al. (2017) demonstrated the efficacy of using the EPOC in the classroom for teaching neuroscience to undergraduate students. Further Dikker et al. (2017) used multiple EPOC systems concurrently to show neural-synchrony across students in a classroom setting. While there is a clear potential for such systems to efficiently gather data in remote locations such as the personal homes of patients, it is necessary to evaluate how data collection in remote contexts that are not electromagnetically-shielded compares with typical research settings. As mentioned above, most EEG recordings take place in electrically-shielded rooms. This is done to attenuate the amount of electrical noise that EEG devices pick up (Jackson & Bolger, 2014), which may be greater than the relatively weak electrical signals output from the brain. Though not all modern research-grade EEG systems use electrical shielding, and its necessity can be questioned (Ledwidge & Ramsay, 2018), whether consumer-grade EEG systems such as the EPOC record research-grade signals without electrical shielding is unknown. Previous research mentioned above has of course used the EPOC outside of electrically shielded environments however, surprisingly, a direct comparison of shielded and unshielded conditions has yet to be performed and would provide added confidence in non-laboratory research programs.

The EPOC uses a driven right leg (DRL) noise-cancellation system, similar to BioSemi EEG systems (www.biosemi.com). While DRL is aimed at attenuating ambient electrical noise via active cancelation, the quality of the system in EPOC is yet to be systematically tested. Such a comparison is critical for determining the reliability of EEG recordings beyond research settings. Establishing that shielding is not required for reliable EEG recording provides assurance for using EPOC outside traditional settings, for example, in classrooms (e.g., de Wit et al., 2017; Dikker et al., 2017).

To address this question, we compared EPOC’s ERP recordings in two settings; an electrically-shielded laboratory and an unshielded quite room. Specifically, we use a similar paradigm to de Lissa et al. (2015), comparing the N170 amplitude and latency to upright and inverted human- and wristwatch-face stimuli. The occipito-temporal N170 (negative brain potential occurring approximately 170 ms after stimulus onset) is reliably larger for faces than other objects (Bentin, Allison, Puce, Perez, & McCarthy, 1996; Eimer, 1998; Rossion & Jacques, 2007). The N170 is also sensitive to face inversion, which delays the timing of the peak and often enhances its amplitude (Itier, Latinus & Taylor, 2006; Itier & Taylor, 2004; Linkenkaer-Hansen et al. 1998; Rossion et al., 2000). The N170 is therefore a good candidate for comparing the effect of recording context with EPOC. However, a common issue in classical statistical analysis is an inability to determine when two things do not significantly differ. Therefore, in the current article, we assessed the degree of difference between the N170 signals obtained in both conditions using Bayesian statistics. This allowed us to quantify to what degree we should favor the null hypothesis (no difference) compared to the alternative hypothesis (difference). In addition, the similarity of the waveforms of ERPs recorded in the two contexts was assessed through intra-class correlations (Shrout & Fleiss, 1979).

By recording the face-sensitivity and face-inversion effects of the N170 in shielded and unshielded contexts, we have found that EPOC records comparable patterns of face-sensitivity and face-inversion effects in both contexts. This was established at both the mean and single-trial level. Further, the intra-class correlation was high, indicating similar waveforms in both contexts. These results suggest that EPOC would be well-suited to record reliable EEG data in at least some contexts outside of typical research settings and may be warranted in such places as schools or doctor’s surgeries.

## 2. Materials and Methods

The Human Ethics Committee at Macquarie University approved the methods used in this study. All participants gave their informed and written consent to participate in the study.

### 2.1. Participants

Our sample size was based on a set timeframe (one month), in which we expected to be able to collect 15 participants. We applied a Bayesian stopping rule to decide whether additional data would be required, however the number of participants proved adequate. Seventeen Macquarie University undergraduate students ended up taking part in the study; however, only 13 (mean age = 23.2 years, SD = 7.9, 18-49, 8 female, 2 left-handed) were used due to hardware issues.^1^ All participants had normal or corrected-to-normal vision. The order of testing of the recording context was counter-balanced, with 7 participating in the Shielded then Unshielded order, and 6 in the Unshielded then Shielded order.

### 2.2. Stimuli

The stimuli used in the current study were identical to those in the study by de Lissa et al. (2015), which consisted of 300 gray-scale images divided into four conditions containing 75 unique images each. The images were either upright (50%) or inverted (50%) wristwatches or emotionally-neutral Caucasian faces, which were cropped within a standard-sized oval frame only showing internal parts of the faces. The four conditions were hence *upright faces, upright watches, inverted faces* and *inverted watches*. The face images were obtained from seven databases: NimStim (Tottenham, et al., 2002), the Karolinska Directed Emotional Faces (KDEF; Lundqvist, et al., 1998), Gur et al. (2002), Computational Vision Archive (courtesy of Caltech), the MIT-CBCL (Weyrauch, et al., 2004), the Ekman and Friesen face set (Ekman & Friesen, 1976), and a set from Kieran Lee and David Perrett of St Andrews University. Wristwatch stimuli were obtained from the University of Kansas Information and Telecommunication Technology Centre database.

Trials began with a white 500-ms fixation cross in the centre of a computer screen against a black background. This was replaced with a 200-ms upright or inverted face or wristwatch, followed by a blank black screen until a response was made. Participants were asked to indicate whether the image was upright or inverted via a binary keyboard response. A response was followed by a 2000-ms “blink” screen before a new trial commenced. The stimuli were presented by Experiment Builder software (version 1.6.1) on a standard MacBook P8600 13-inch laptop at a distance of 40 cm from the participant. As such, each image was 13.1^0^ × 10.4^0^ degrees of visual angle.

### 2.3. Contexts

Two contexts were used for EEG testing; an electrically-shielded research laboratory used for electrophysiological experiments, and an unshielded room. The unshielded room consisted of a chair and a desk in a dimmed room without electrical shielding. There was a window to the outside, with blinds drawn. Both rooms were comparable in size and brightness. The levels of ambient noise were consistent, and there were no electrical devices being run in either room apart from those used for recording.

### 2.4. Emotiv EPOC EEG system

The Emotiv EPOC EEG system (original model, purchased 2010) uses 16 gold-plated sensors (coated in an electrochemically-active material infused polymer) at locations corresponding to AF3, AF4, F3, F4, FC5, FC6, F7, F8, T7, T8, P7, P8, O1, O2, Common Mode Sense (CMS, left earlobe), and Driven Right Leg (DRL, right earlobe) on the international 10-20 system. The CMS and DRL provide an active reference system. The sensors were connected to the participants’ scalps through felt pads soaked in saline solution, along with Signa Gel electrode gel between the felt pads and the participant’s scalp.

EEG signals detected by the EPOC electrodes were pre-processed within the headset (programmed by the manufacturer), passing through a 0.16-Hz high-pass filter pre-amplification process, as well as an 83-Hz low-pass filter, before being digitized at 2048 Hz. Two notch filters were then applied at 50 and 60 Hz before further low-pass filtering was performed with a 43-Hz cut-off. The signal was then down-sampled to 128 Hz and transmitted to a recording computer via a proprietary wireless signal. The EPOC system uses a proprietary impedance value system.

The Emotiv EPOC EEG device was modified for the current study in two ways. (1) The marker-triggering circuit involved injecting electrical signals into two EEG channels (T7 and T8) at the onset of a stimulus (Badcock et al., 2013; de Lissa et al., 2015; Thie, 2013). These signals were used to create epochs for ERP analysis. The marker system used here was identical to that used in previous work (for technical details, see de Lissa et al., 2015). Triggers were sent via a phototransistor which was activated by a white square presented in the corner of the screen with each stimulus. This white square was occluded by the phototransistor.

(2) The online reference system EPOC was modified to use the left and right earlobes as CMS (Common Mode Sense) and DRL (Direct Right leg) locations, respectively. This was done to avoid using the standard EPOC reference systems of either the left mastoid or left P3 electrode areas, which are not optimal locations when recording occipito-temporal N170 peaks due to their proximity to the epicentres of this activity as measured from the scalp (Joyce & Rossion, 2005). The modifications were achieved by re-routing the left (CMS) and right (DRL) sensors of the EPOC system to the participants’ left and right earlobes by wiring shortened (approximately 5 cm) Ag-AgCl EasyCap electrodes into the fixed M1 and M2 sensors (for further details, see de Lissa et al., 2015).

### 2.5. EEG offline processing

The EEG data was processed offline with EEGLAB version 14.1.2 software (Delorme & Makeig, 2004). The EEG data incorporated the left earlobe as an online common-mode sense reference point online and was filtered through a band-pass of 0.1– 30 Hz with a 12 dB/octave roll-off. EEG artefacts were excluded by visual analysis and then ocular artefact correction was performed through independent component analysis (ICA; Viggario, 1997; Delorme & Makeig, 2004). Epochs were created (−102 – 602 ms, relative to stimulus onset) and baseline-corrected to the 102 ms period preceding stimulus onset. Epochs containing EEG signals exceeding +/−150 µV were excluded from further analysis. The accepted epochs for each participant were averaged to produce ERPs for P7 and P8 for each condition and context.

Amplitudes, latencies, the number of accepted epochs, and the signal to noise ratio (SNR) were analyzed. Amplitudes were taken as the average activation surrounding the N170, 125 to 195 ms following stimulus onset. N170 latencies were taken as the timepoint of minimum activation within the same time-period. The number of accepted epochs is those included in the average waveforms after epochs with extreme values were removed, or those missing due to misalignment of the infrared event-marking system. The SNR was calculated as the root mean squared (RMS) of the signal from 102 ms to 203 ms, divided by the standard deviation of the signal from −102 ms to 0 ms, on each trial separately at the P7 and P8 electrodes (see Maidhof, Rieger, Prinz, & Koelsch, 2009; Marco-Pallares, Cucurell, Münte, Strien, & Rodriguez-Fornells, 2011).

In addition to the average activations, we also calculated the N170 amplitude on each individual trial at the P7 and P8 electrodes, which were then used for linear mixed effects (LME) modelling, see next section. Further, the SNR was calculated on a single-trial basis as per for the mean waveforms.

### 2.6. Data analysis

Intra-class correlations (ICCs) were also included as a descriptor of the shape and amplitude of the waveforms between contexts (as in previous work: Badcock et al., 2013; 2015; Bishop & McArthur, 2005; de Lissa et al., 2015; McArthur & Bishop, 2005).

There were three sets of inferential tests on the averaged data. The first was to confirm the N170 effects. This included testing (1) whether faces produced greater N170 amplitudes than wristwatches; (2) whether faces, but not wristwatches, produced greater N170 amplitudes for upright versus inverted images; and (3) whether faces, but not wristwatches, produced longer N70 latencies for upright versus inverted images.

The second tested whether context (i.e., recording in shielded versus unshielded room) affected N170 amplitude, latency, or the number of accepted epochs. All inferential tests included specific main effects and/or interactions (rather than an omnibus set of comparisons) and we report the standard significance test of analyses of variance (ANOVA) as well as Bayesian ANOVAs. Bayesian statistics were included to confirm any null effects; i.e., a Bayes Factor less than 0.3. Bayes Factors greater than 3 were treated as evidence of differences. The analyses were conducted in RStudio (R Core Team, 2019; RStudio Team, 2015) and the BayesFactor package (Version 0.9.12-4.2, Morey & Rouder, 2018), using the default settings.

The final test on the averaged data was a comparison between the SNR under the shielded and unshielded contexts. This was done using a classical paired t-test, using R, and a Bayesian paired t-test using the BayesFactor package.

In addition to the statistical tests on the averaged data, we built an LME on single-trial N170 amplitude data and the SNR. Starting from the null model which only included subjects as a random effect, fixed effects were added sequentially, including interactions, and only kept if the model fit was significantly improved, as indicated by a significant *Χ*^*2*^ result and a decrease in the Bayesian Information Criterion (BIC). If multiple factors would result in a significant effect, the factor which decreased the BIC the most was kept.

Amplitude and SNR were used as dependent factors, while stimulus type, stimulus orientation, the orientation of the stimuli, hemisphere (P7/P8), and recording context were successively added as independent factors. The subjects were included as random effects. Models were fit using the lme4 (Bates et al., 2015) package in R, and the significance of each effect in the winning model was calculated using the lmertest package (Kuznetsova, Brockhoff, & Christensen, 2017).

## 3. Results

Average ERP waveforms for the left and right hemispheres (i.e., P7 vs. P8) by stimulus type (i.e., faces vs. watches) by stimulus orientation (i.e., upright vs. inverted) by recording context (i.e., shielded vs. unshielded room) are presented in Figure 1. Key waveform differences (i.e., Faces vs. Wristwatches and Upright vs. Inverted Faces) are presented in Figure 2. Column scatter plots for the key variables of interest for amplitude and latency are reported in Figure 3 and Figure 4 respectively. The amplitude and latency patterns by stimulus type and orientation were consistent with the background literature on the N170 face effect. The intra-class correlations (ICCs) comparing the waveforms in shielded and unshielded contexts were medium in strength and all significant based on the confidence intervals not overlapping with zero (see Table 1).

**Table 1.** Mean Intra-class correlations comparing waveforms recording in the Shielded and Unshielded context.

| Electrode | Stimulus | Orientation | Mean | Lower CI | Upper CI |
| --- | --- | --- | --- | --- | --- |
| P7 | Faces | Upright | 0.64 | 0.54 | 0.75 |
|  |  | Inverted | 0.70 | 0.59 | 0.81 |
|  | Watches | Upright | 0.65 | 0.48 | 0.81 |
|  |  | Inverted | 0.68 | 0.57 | 0.79 |
| P8 | Faces | Upright | 0.51 | 0.30 | 0.71 |
|  |  | Inverted | 0.70 | 0.56 | 0.84 |
|  | Watches | Upright | 0.65 | 0.54 | 0.77 |
|  |  | Inverted | 0.69 | 0.58 | 0.79 |

**Figure 1.**
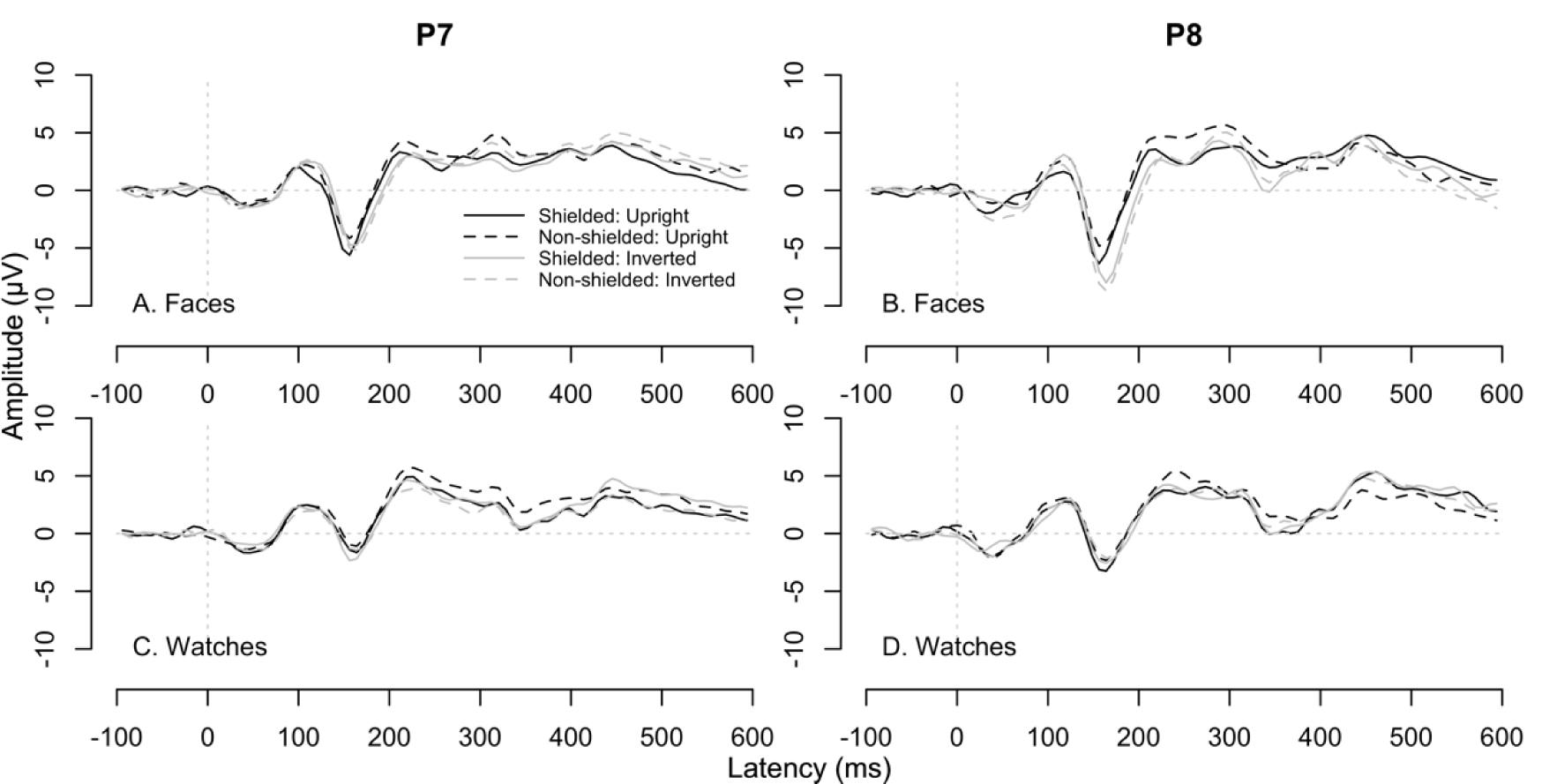
Event-related potential waveforms for hemisphere, stimulus type, stimulus orientation, and recording context.

**Figure 2.**
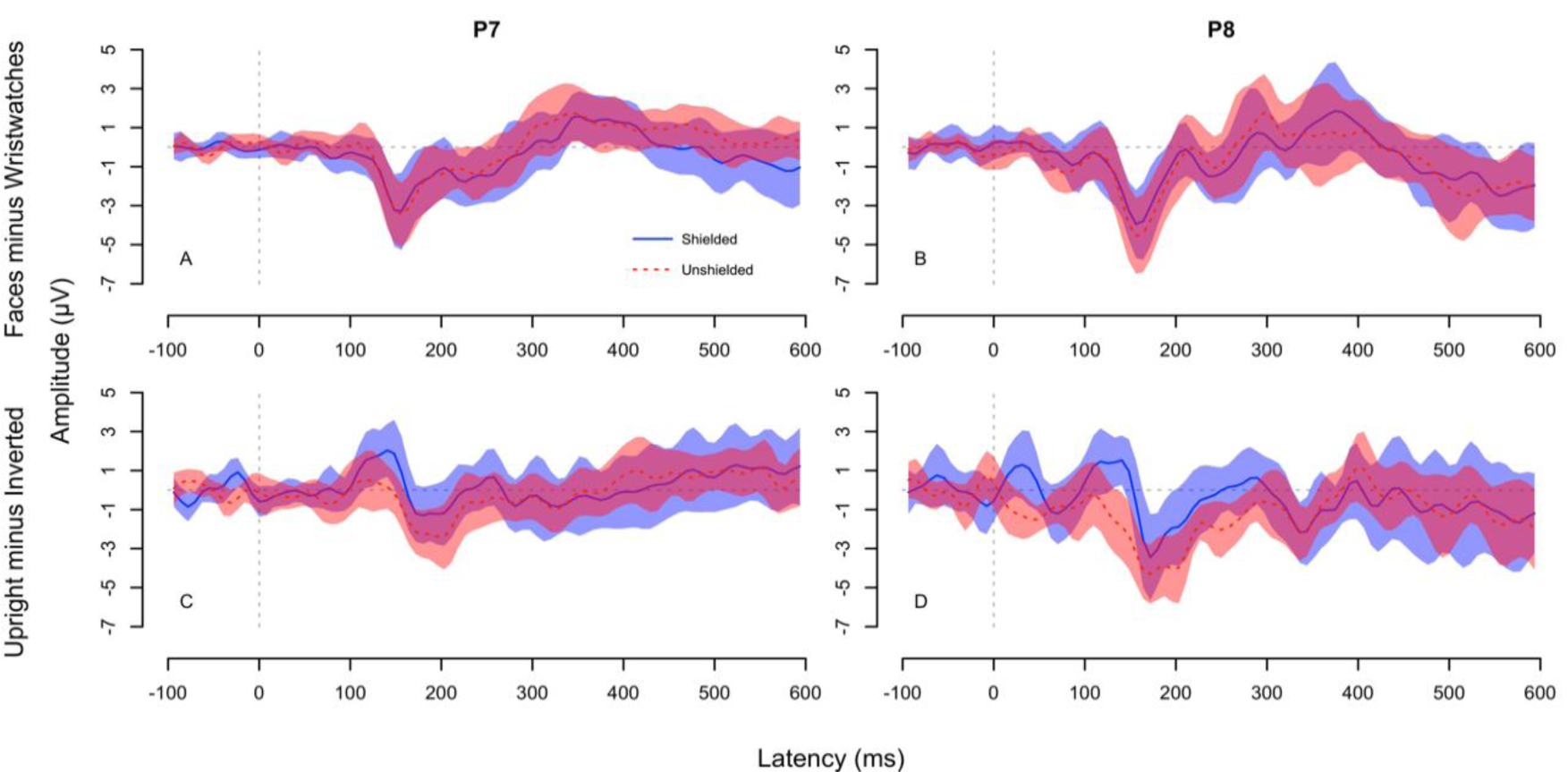
Event-related potential waveform differences for Faces minus Wristwatches (top row) and Upright minus Inverted faces (bottom row). Shaded areas represent the 95% confidence intervals.

**Figure 3.**
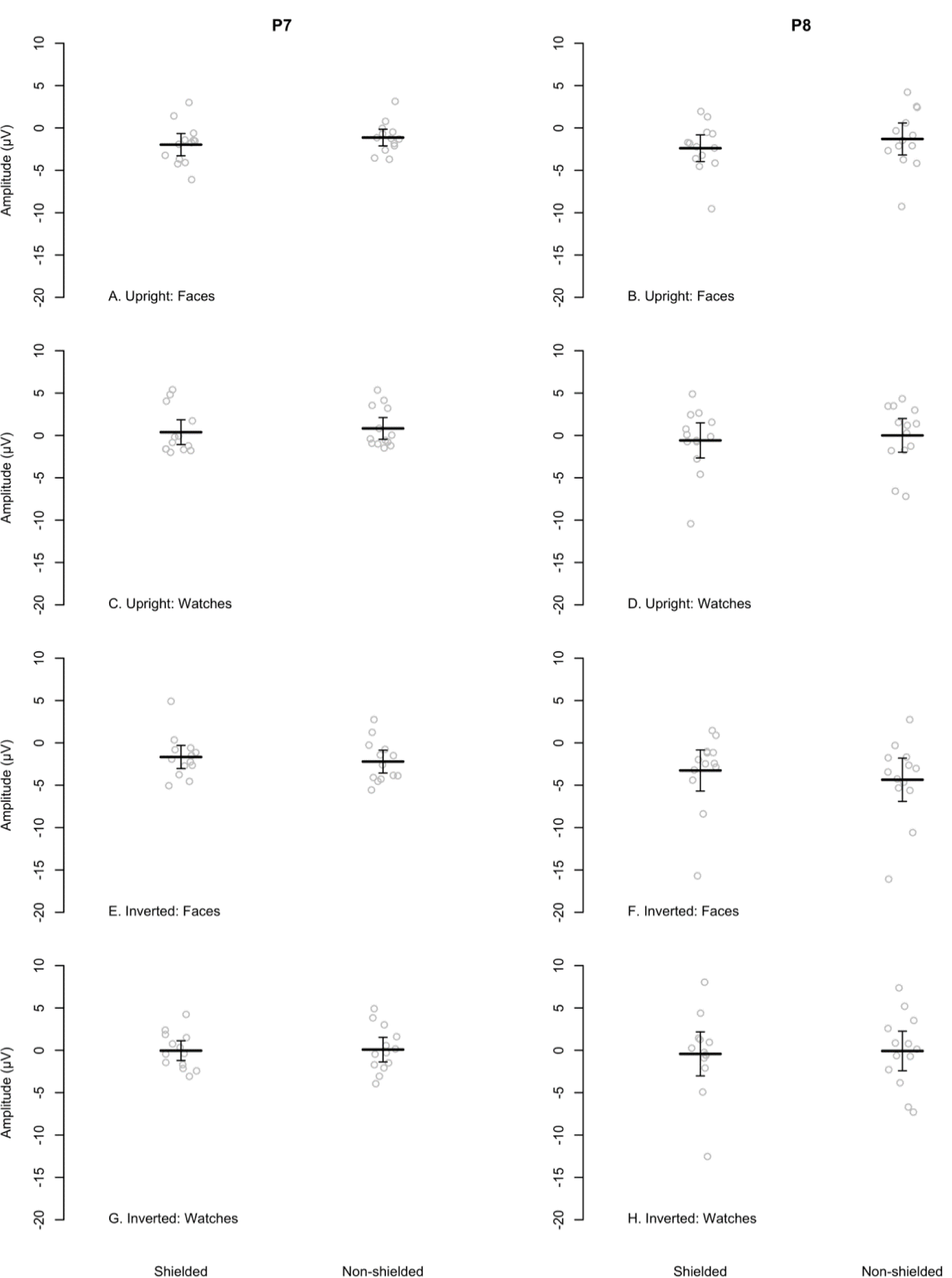
Column scatter plots of the mean activation amplitude for the N170 (125 to 195 ms following stimulus onset). Group and individual means are presented and error bars represent the 95% confidence intervals for the group.

**Figure 4.**
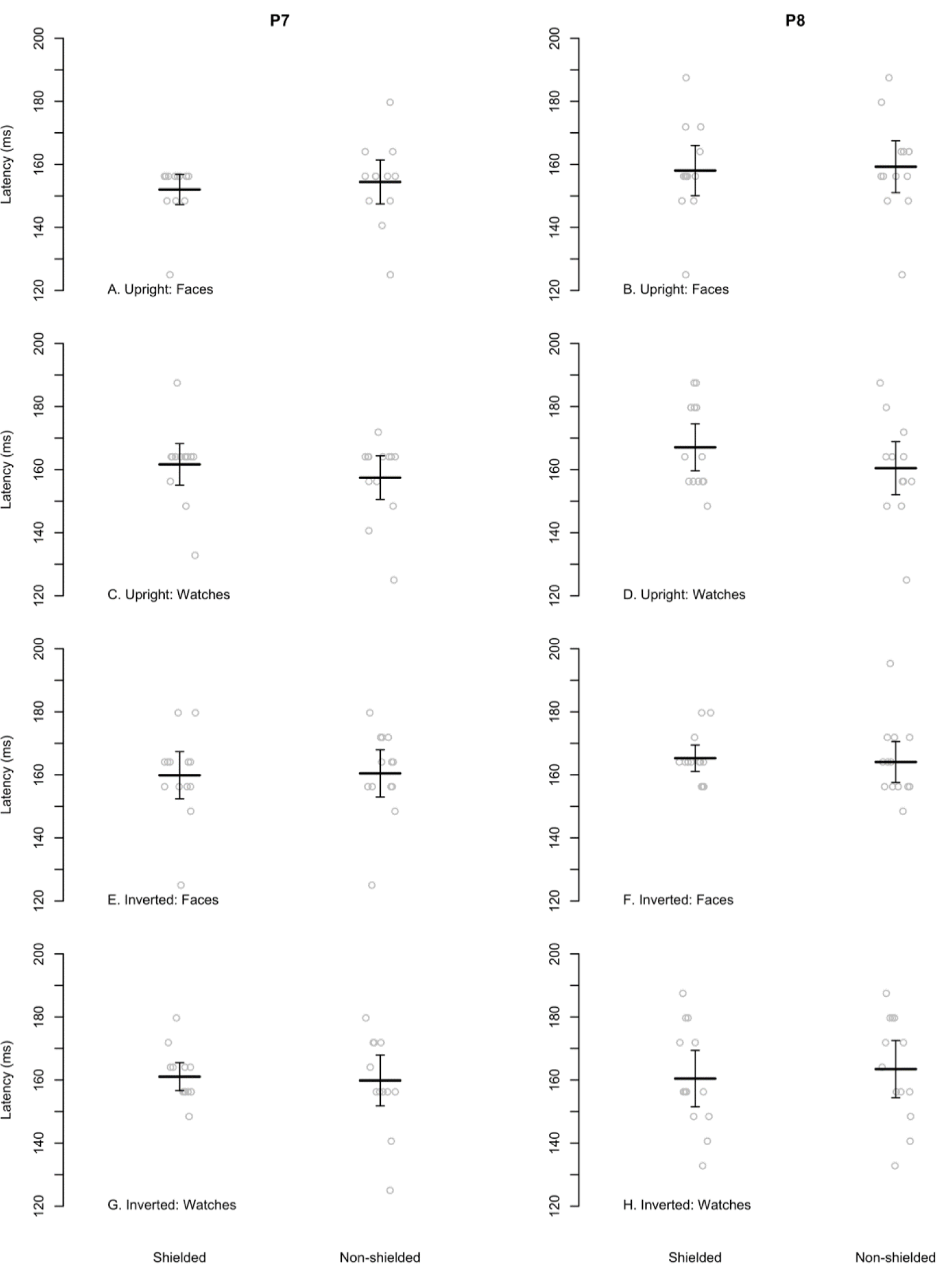
Column scatter plots of the latency of the minimum amplitude between 125 to 195 ms following stimulus onset (i.e., the N170). Group and individual means are presented and error bars represent the 95% confidence intervals for the group.

### 3.1. Discarded epochs

Context did not affect the number of EEG epochs rejected in the artefact-rejection processe (detailed in Section 2.5); F(1, 12) = 2.45, p = 0.12, *η*_p_^2^ = 0.01, BF_10_ = 0.52 (note: this is an inclusive BF_10_ but, given the weak effect size, the practical impact of any difference is negligible). The means of accepted epochs were therefore comparable between contexts: 54.90 (SD = 7.62, min = 32, max = 60) epochs were retained in the shielded context, 55.69 (SD = 4.73, min = 42, max = 60) in the unshielded context.

### 3.2. Amplitude

Faces produced higher amplitude responses than watches; F(1, 12) = 42.36, p < .001, *η*_p_^2^ = 0.18, BF_10_ = 1.44^7^. Stimulus type interacted with orientation (i.e., upright vs. inverted); F(3, 12) = 16.36, p < .001, *η*_p_^2^ = 0.2, BF_10_ = 6.38^6^; and was significant for faces (inverted larger negative peak than upright); F(1, 12) = 5.07, p < .05, *η*_p_^2^ = 0.05, BF_10_ = 1.91; but not for watches; F(1, 12) = 0.42, p = 0.52, *η*_p_^2^ = 0, BF_10_ = 0.24. There was no effect of Context on N170 amplitude; F(1, 12) = 0.33, p = 0.57, *η*_p_^2^ = 0, BF_10_ = 0.18.

### 3.3. Latency

For minimum peak latency, the stimulus type interacted with orientation (i.e., upright vs. inverted); F(3, 12) = 4.9, p < .001, *η*_p_^2^ = 0.07, BF_10_ = 3.7; being significant for faces (i.e., longer latency for inverted faces); F(1, 12) = 15.23, p < .001, *η*_p_^2^ = 0.14, BF_10_ = 127.83; and again, not for watches; F(1, 12) = 0.05, p = 0.82, *η*_p_^2^ = 0, BF_10_ = 0.21. There was no effect of Context on N170 latency; F(1, 12) = 0.3, p = 0.58, *η*_p_^2^ = 0, BF_10_ = 0.17.

### 3.4 SNR

The SNR was not significantly affected by context: shielded (M = 5.08, SD = 0.65) vs. unshielded (M = 5.35, SD = 0.57), t(12) = .592, p = 0.565, *d* = 0.14, BF_10_ = 0.32. The BF provides moderate evidence against an actual difference in SNR between the two contexts.

### 3.5 Single-trial Amplitude LME

Compared to the null model, the addition of hemisphere (i.e., electrodes) reduced the BIC, *Χ*^*2*^(1, 4) = 178.7, *p* < .001, ΔBIC = −169. In the next step, the addition stimulus type maximally decreased BIC, *Χ*^*2*^(2, 6) = 89.50, *p* < .001, ΔBIC = −71. From this model, neither the addition of context, *Χ*^*2*^(4, 10) = 8.36, *p* = .079, ΔBIC = +30, nor the addition of stimulus orientation, *Χ*^*2*^(4, 10) = 9.37, *p* = .052, ΔBIC = +29, improved model fit, and thus were not included.

The final model showed that the mean amplitude of the N170 was lower (i.e., larger) following the presentation of faces (−11.65 µV) compared to watches (−9.00 µV, p < .001). Further, the N170 was larger in the P8 electrode (−12.11 µV) compared to the P7 electrode (−8.52 µV, p < .001). The interaction between these two factors was not significant (p = .601).

### 3.6 Single-trial SNR LME

For SNR, only the addition of context resulted in a significant *Χ*^*2*^ effect, however this failed to improve the BIC, *Χ*^*2*^(1, 4) = 8.07, *p* = .004, ΔBIC = +2. Therefore, no model appeared to parsimoniously improve the model fit beyond the null model. However, because the context in which the recording occurred was of primary concern to the article at hand, we examined the results of this model. It showed that the SNR was higher in the non-shielded context (M = 2.01, SD = 0.66) rather than in the shielded context (M = 1.91, SD = 0.58).

## 4. Discussion

In the current article we asked whether electrical shielding significantly affects the electrophysiological recordings provided by Emotiv’s EPOC EEG system. To answer this question, participants performed the same task twice, once in a shielded room and once in an unshielded room. Participants were required to observe upright and inverted, human- and wristwatch-faces, a task previously used to demonstrate that EPOC provides EEG recordings comparable to a research-grade system in shielded laboratories (de Lissa et al., 2015).

Results indicate that classic face-sensitive N170 effects were found in electrodes P7 and P8; faces resulted in larger amplitude N170 responses, and inverted faces resulted in longer latencies than upright faces (e.g., Bentin et al., 1996; Eimer, 1998; Itier et al., 2006, Itier & Taylor, 2004; Linkenkaer-Hansen et al., 1998; Rossion et al., 2000). However, more relevant to our research question, the findings demonstrated that the EPOC provides comparable EEG recordings in shielded and unshielded environments. Firstly, we showed that the overall waveforms in each context were highly correlated; the data provided by the EPOC followed the same pattern whether recorded in a shielded or unshielded environment. Further, comparing the face-sensitive N170 in both contexts showed no significant difference in either amplitude or latency. Consistently, Bayesian analyses provided moderate support (in accordance with the Jeffreys, 1961, Bayes factor interpretation) for no difference.^2^ Finally, the Bayesian analysis also provided moderate support against an actual difference in SNR between the shielded and unshielded contexts.

Analyzing the data at a single trial level, we found an effect of stimulus type (i.e., faces > watches), and similarly, a larger effect at electrode P8 compared to P7. However, the single trial analysis did not provide evidence for a difference in N170 based on whether shielding was present. Taken together, the evidence appears to show that the EPOC and its DRL noise-cancellation system, is appropriate for use outside a shielded research laboratory, at least in a quiet room.

This finding has ramifications for both the interpretation of prior research, and the acceptability of unorthodox research environments. As mentioned in the introduction, EPOC has previously been used to record EEG while participants walk outside (Debener et al., 2012; De Vos et al., 2014). While this is an important demonstration, it is useful to know that these results are most likely equivalent to those gained in a research laboratory content. Further, knowing that the recording achieved in an unshielded room is comparable to in a laboratory setting is useful for those who wish to employ EEG in learning environments (e.g., de Wit et al., 2017; Dikker et al., 2017).

An additional important point is that there are many barriers to entry for would-be scientists; specialty knowledge, time, access to resources, to name a few. However, research such as that conducted here can serve as a small step to remove the hold institutions traditionally have on ‘doing science.’ Showing that EEG can be comparably recorded outside the laboratory with a relatively small investment is an important step in the democratization of science.

While these results are promising, it is worth noting some relevant future extensions to the current research. Firstly, we have clarified that the EPOC system is suitable for use outside of electrical shielding. This aligns with previously work (e.g., Debner et al., 2012; De Vos et al., 2014), but adds a direct control comparison of a shielded environment. While the Emotiv systems are currently popular for research purposes, other brands of consumer-grade EEG have yet to be tested. Secondly, our unshielded environment was relatively conducive to research. How EPOC would fare in a busier environment is unknown. While the current research seems to indicate the recording would be of a comparable quality to a shielded room, how the general noise (electrical or otherwise) around the recording device affects the recording is not directly tested. As another further direction for this research, it would be interesting to systematically vary the noise injected into the headset in multiple locations, such as shielded rooms, outdoors or classrooms, and examine whether the variance of this noise is similar across locations.

## 5. Conclusion

In the current article, we compared the performance of the Emotiv EPOC consumer-grade EEG system in shielded and unshielded recording contexts. Analysis of both the N170 face-sensitive event-related potentials showed that EPOC performs comparably in both contexts. This provides empirical evidence validating the use of the EPOC in the unshielded setting and allows for research-backed comparisons between research conducted using this device in laboratory and non-laboratory settings.

## Acknowledgements

This research was funded by the Australian Research Council Centre of Excellence in Cognition and its Disorders (CE110001021). All authors conceived the design of the project. PD programmed the stimulus presentation. JW & SS collected the data. JW & PD processed the EEG files. NAB analysed the data and created the figures. All authors contributed to writing the manuscript.

## Footnotes

1 For one participant, EEG was not recorded. For the remaining three, the trigger system failed and thus data could not be timelocked. Trigger failure occurred when either the connection to the custom-made trigger device (see below) was lost, for example if the wires lost contact with the selected trigger channels, or if the device lost power during the session, due to the battery running out.

2 It is worth noting that Bayes Factor was inconclusive in regards to the number of epochs discarded given a shielded or unshielded context, though the difference was negligible from a practical perspective.

